# mRNP architecture in translating and stress conditions reveals an ordered pathway of mRNP compaction

**DOI:** 10.1101/366690

**Authors:** Anthony Khong, Roy Parker

## Abstract

Stress granules (SGs) are non-translating mRNP assemblies that form during stress. Herein, we use multiple smFISH probes for specific mRNAs to examine their SG recruitment and spatial organization. We observed that ribosome run-off is required for SG entry with long ORF mRNAs being delayed in SG accumulation, revealing SG transcriptome changes over time. Moreover, mRNAs are ~20X compacted from an expected linear length when translating and compact ~2 fold further in a stepwise manner beginning at the 5’ end during ribosome run-off. Surprisingly, the 5’ and 3’ ends of the examined mRNAs were separated in non-stress conditions, but in non-translating conditions, the ends of AHNAK and DYNC1H1 mRNAs become close, suggesting the closed-loop model of mRNPs preferentially forms on non-translating mRNAs. These results suggest translation inhibition triggers a mRNP reorganization that brings ends closer, which has implications for the regulation of mRNA stability and translation by 3’ UTR elements and the poly(A) tail.

## INTRODUCTION

Stress granules (SGs) are transient membraneless organelles of non-translating mRNA-protein complexes (RNPs) that form when translation is limited (Buchan and Parker, 2009; Panas et al., 2016; Protter and Parker, 2016). SGs are important because they are a cellular marker for translation status, play a role in the stress response (Kedersha et al., 2013), and mutations that inhibit SG disassembly or clearance are implicated in several degenerative diseases such as amyotrophic lateral sclerosis (ALS) and multisystem proteinopathy (Buchan et al., 2013; Dewey et al., 2012; Kim et al., 2013; Li et al., 2013; Mackenzie et al., 2017; Ramaswami et al., 2013). Moreover, the study of SGs may provide new insights into the assembly, organization, and functions of other non-membrane bound RNA bodies such as the nucleolus, Cajal bodies, paraspeckles, and processing bodies.

SGs are enriched for mRNAs that are long and poorly translated (Khong et al., 2017; Namkoong et al., 2018). This suggests a model wherein long mRNPs that exit translation during stress form interactions with other long non-translating mRNPs leading to the formation of SGs. Some interactions between mRNAs that promote SG formation are between mRNA binding proteins that are thought to provide cross-links between individual mRNAs and thereby enhance SG assembly (reviewed in Protter and Parker, 2016). However, intermolecular RNA-RNA interactions can contribute to SG formation and to defining the SG transcriptome, which is suggested by the observation that self-assembly of RNA *in vitro* can largely recapitulate the yeast SG transcriptome (Van Treeck et al., 2018). An unresolved issue is the relative timing of mRNAs exiting translation, how translation affects the organization of the mRNP, and the timing of mRNAs accumulating in SG.

The timing of SG formation and the enrichment of long mRNAs in SGs creates a conundrum. This is because SGs form within the first 10-15 minutes after the addition of arsenite (Kedersha et al., 2000; Wheeler et al., 2016), yet mRNAs with long open reading frame (ORF) such as the SG-enriched mRNA AHNAK and DYNC1H1 (ORF > 10 kb) (Khong et al., 2017) require at least 15 minutes for ribosome run-off once translation initiation is blocked. One possibility is that these long mRNAs can accumulate in SGs once a portion of their ORF is exposed and devoid of ribosomes, even if ribosomes near the 3’ end of the ORF are still elongating. Another possibility is that elongating ribosomes are removed from these mRNAs without having to reach the termination, perhaps by a mechanism analogous to ribosome quality control (Brandman and Hegde, 2016; Brandman et al., 2012; Chiabudini et al., 2014; Harigaya and Parker, 2010; Shoemaker et al., 2010; Shoemaker and Green, 2012). Finally, it is also possible that mRNAs with long ORF are slower at getting to SGs, which would require the SG transcriptome to change over time.

To examine how mRNAs exit translation and enter SGs, we used multiple smFISH probes for specific mRNAs to examine the timing of when those mRNAs enter SGs, and their spatial organization, which revealed key aspects of mRNA targeting to SGs. First, complete ribosome run-off is required for mRNAs to enter SGs with mRNAs with long ORFs being delayed in SG accumulation. This demonstrates that SG transcriptome changes over time. We also observed that mRNAs are compacted from an expected linear length when translating, and compact even further in a step-wise manner due to ribosome run-off. We do not see evidence for the closed loop model of mRNP organization with the mRNAs examined while they are engaged in translation, although the distance between the 5’ and 3’ ends of long mRNAs shrinks disproportionally compared to the rest of the mRNAs when mRNAs are untranslated. We suggest the possibility that the closed loop structure of mRNPs preferentially forms on non-translating mRNPs.

## RESULTS

### mRNAs with long ORF are recruited slower to SGs than mRNAs with shorter ORF

To determine the relationship between SG assembly and the recruitment of mRNAs with long ORF, we measured when several SG-enriched mRNAs (Khong et al., 2017) with various ORF lengths were recruited to SGs in cells treated with arsenite for 15’, 30’, 45’, and 60’ by smFISH. These include AHNAK, DYNC1H1, NORAD, PEG3, ZNF704, CDK6, and NORAD RNAs. AHNAK and DYNC1H1 mRNAs have long ORF (~17.5kb and 14kb respectively), while the PEG3 and ZNF704 mRNAs have shorter ORF (~4.7kb, 1.2kb respectively). The CDK6 mRNA is valuable since it has a short ORF (~1 kb), but has a very long 3’ UTR (~ 10 kb), allowing us to distinguish effects of the overall transcript length from ORF length. We also examined when a lincRNA, NORAD, is recruited to SGs. The predicted ribosome run-off times for these mRNAs once translation initiation is blocked are shown in Table 1. We performed these experiments in U-2 OS cells, where arsenite induces robust eIF2α phosphorylation, an approximate marker for when translation initiation is inhibited, at ~8’ (Wheeler et al., 2016).

**Table 1.** Predicated ribosome run-off time

| mRNA | total length<br>(nt) | CDS length<br>(nt) | Predicted ribosome run<br>-off time (18nt/sec) |
| --- | --- | --- | --- |
| AHNAK | 18,836 | 17,673 | ~16 min |
| DYNC1H1 | 14,361 | 13,941 | ~13 min |
| PEG3 | 8,765 | 4,767 | ~4 min |
| ZNF704 | 14,403 | 1,239 | ~1 min |
| CDK6 | 11,661 | 981 | ~1 min |

A key result was that individual RNAs accumulated in SGs at different times in a manner correlated with the length of their ORF. Specifically, we observed that when cells were stressed for 30 minutes with NaAsO_2_, the AHNAK and DYNC1H1 mRNAs with long ORFs were minimally recruited to SGs (12%) (Figure 1A, B, Supplemental Figure 1). In contrast, RNAs with shorter ORF or no ORF were recruited to a greater degree (39-55%) at 30 minutes (Figure 1A, B, Supplemental Figure 1). At 60 minutes, all the examined SG-enriched RNAs have reached their maximal level of enrichment in SGs (Figure 1A, B, Supplemental Figure 1). These results suggest the mRNA composition of SGs changes over time during arsenite stress and although mRNAs with longer ORF are highly enriched in SGs (Khong et al., 2017), they accumulate slower.

**Figure 1.**
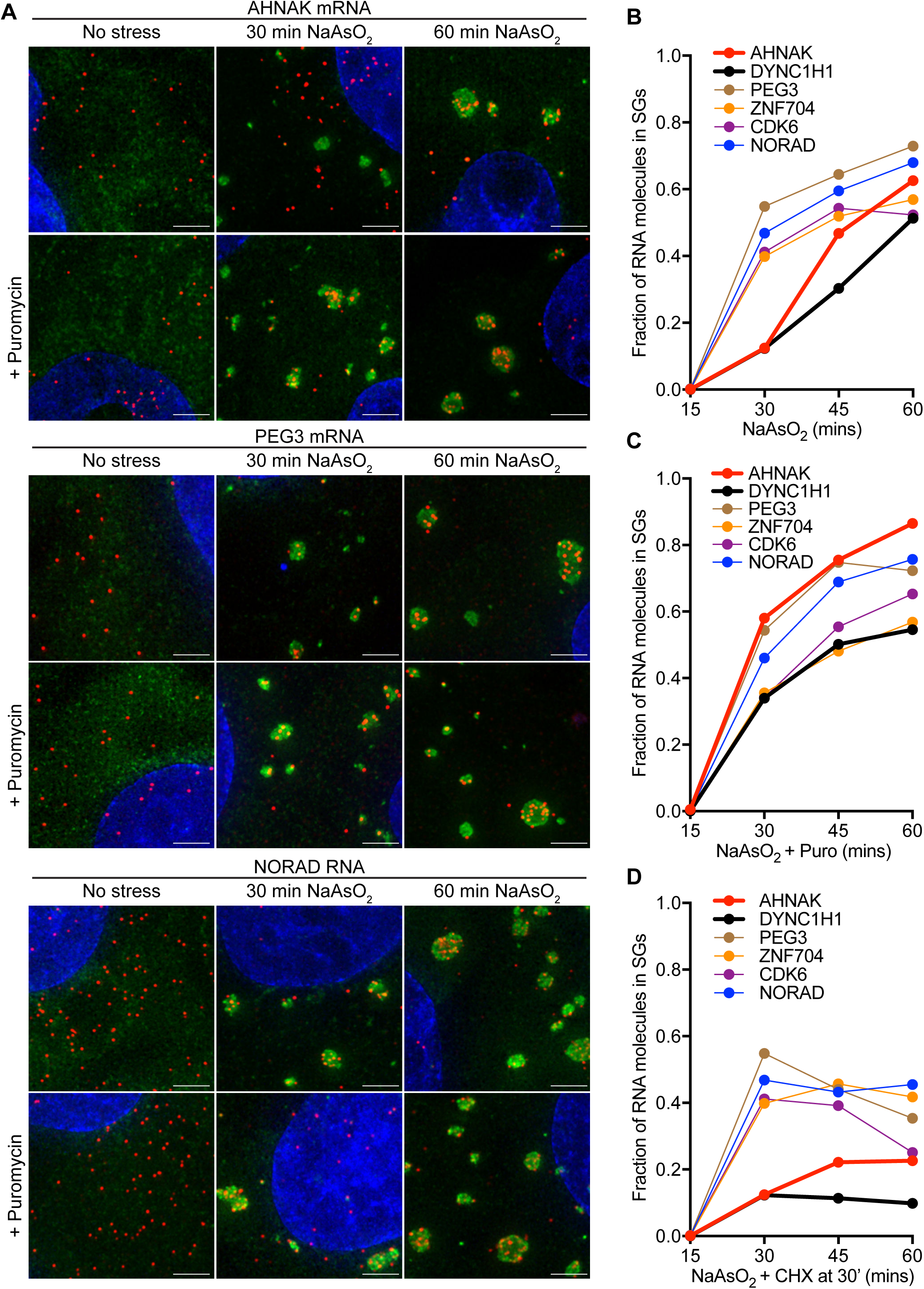
mRNA recruitment to SGs is dependent on when ribosomes run-off elongation after translation inhibition. **(A)** Representative smFISH images acquired for three different transcripts (AHNAK, DYNC1H1, and NORAD) for U-2 OS cells treated with 0.5 mM NaAsO_2_ with or without 10 μg/mL puromycin for 30’ or 60’. Cells were stained with nuclei (blue), G3BP1 antibody (green), and specific transcripts by smFISH (red). Scale bar: 1 μm **(B)** Fraction of specific RNA molecules found in SGs in U-2 OS cells stressed with 0.5 mM NaAsO_2_ for 15’, 30’, 45’, and 60’. **(C)** Fraction of specific RNA molecules found in SGs in U-2 OS cells stressed with 0.5 mM NaAsO_2_ and 10 μg/mL puromycin for 15’, 30’, 45’, and 60’. **(D)** Fraction of specific RNA molecules found in SGs in U-2 OS cells stressed with 0.5 mM NaAsO_2_ for 15’, 30’, 45’, and 60’. 50 μg/mL cycloheximide was added after cells were stressed for 30’. More than 500 RNAs were counted for each sample.

Two additional observations suggest the difference in the timing of mRNA recruitment to SGs is due to elongating ribosomes. First, treatment of U-2 OS cells with arsenite and puromycin, which releases all elongating ribosomes from mRNAs, causes the AHNAK and DYNC1H1 mRNAs to be recruited to SGs at earlier times (Figure 1A, C, Supplemental Figure 2). Second, treatment of U-2 OS cells after 30 minutes of arsenite exposure with cycloheximide, which traps elongating ribosomes on mRNAs, stops the accumulation of all RNAs in SGs (Figure 1A, D, Supplemental Figure 3).

### AHNAK and DYNC1H1 mRNPs are generally compact under non-stress conditions

In other work, we had observed that during stress the 5’ and 3’ ends of the AHNAK mRNA were often close together (Moon et al., 2018). Similar compaction of three other mRNAs under a variety of stress conditions have been observed (Srivathsan et al., 2018). To examine the overall architecture of mRNAs during normal and stress conditions, and how it related to mRNA entry into SG, we utilized smFISH probes to the 5’ end, the 3’ end, and throughout the middle of the AHNAK and DYNC1H1 mRNAs (Figure 2A, Supplemental Figure 4A). We first used these probes on unstressed cells where mRNAs are engaged in translation. We measured the distances between the center of the signal for each probe in three dimensions (see Methods), which allowed us to determine the distribution of spacing for these probe sets on individual mRNA molecules.

**Figure 2.**
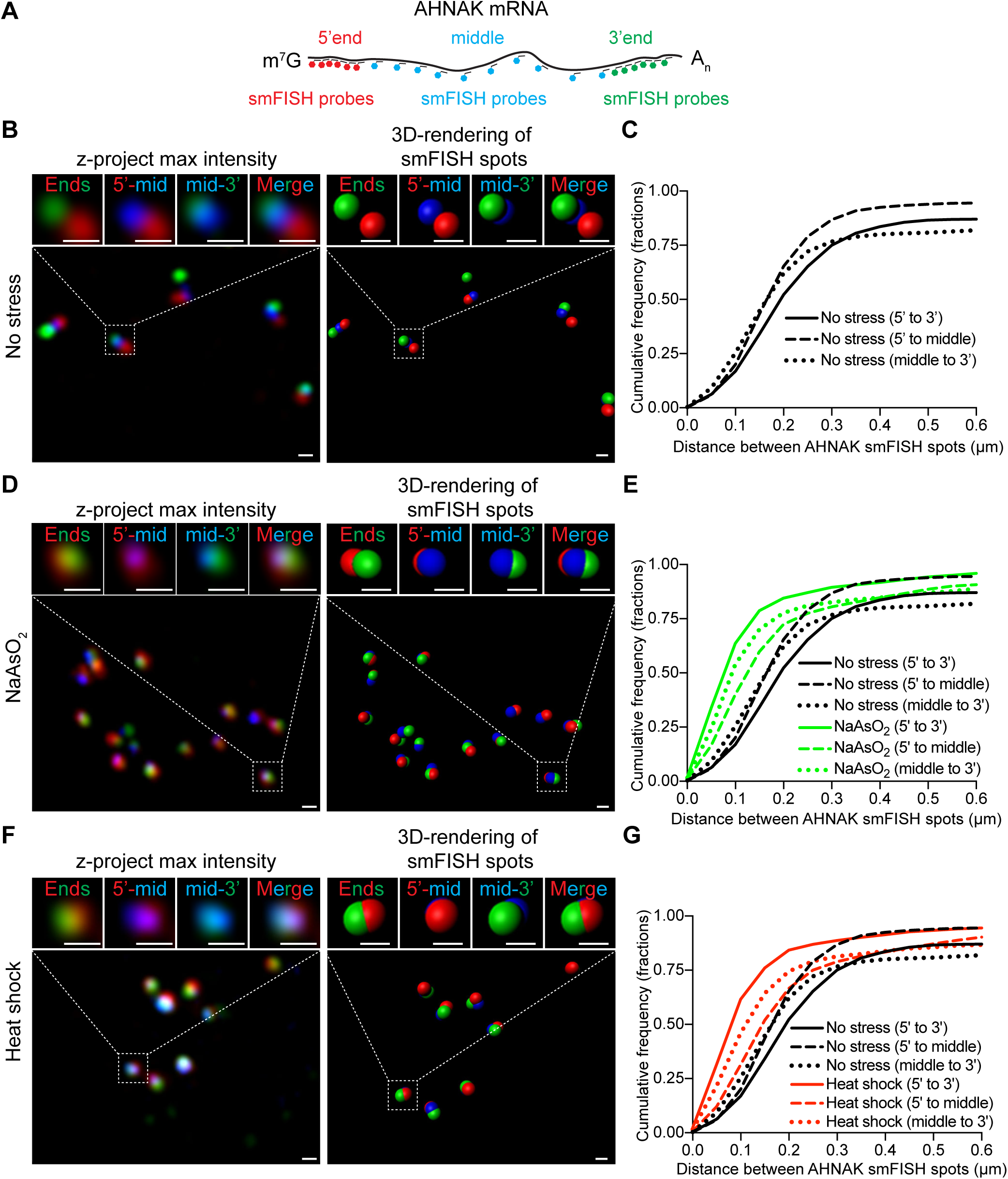
AHNAK mRNPs organization in non-stress and stress conditions. **(A)** Cartoon schematic indicating where smFISH probes bind to AHNAK mRNAs. smFISH probes binding to the 5’ends, middle or 3’ends are labeled with distinct fluorophores and are false-colored red, blue, and green respectively. **(B, D, and F)** Left panels. Representative AHNAK smFISH images of U-2 OS cells that were **(B)** not stressed or **(D)** stressed with 0.500 mM NaAsO_2_ for 60’ or **(F)** heat shock at 42°C for 60’. U-2 OS cells were stained with AHNAK smFISH probes that bind specifically to the 5’ end (false-colored red), middle (false-colored blue), and 3’ end (false-colored green). Right panels. 3D rendering of smFISH spots by Bitplane Imaris imaging analysis software. Scale bar: 250 nm. **(C, E, and G)** Cumulative frequency graphs (in fractions) of smallest distances between 5’ to 3’ end smFISH spots (solid lines), 5’ end to middle smFISH spots (dash lines), and middle to 3’end smFISH spots (dotted lines) in unstressed cells (black), 0.500 mM NaAsO_2_-treated cells (green), and heat shock cells (red). More than 1000 smallest distances were quantified for each sample.

We discovered both AHNAK and DYNC1H1 mRNAs are more compact than expected from linear or hairpin models of translating mRNPs (Figure 2B, Supplemental Figure 4B) with most distances between different segments of AHNAK mRNPs being less than 300 nm (Figure 2B, C). The distances between the 5’ and 3’ ends of AHNAK mRNAs are usually larger (median ~ 200 nm) than the distances between 5’end and middle (median ~ 150 nm) or the 3’ end and middle of AHNAK mRNPs (median ~ 150 nm). These distance measurements are much shorter than expected from the AHNAK mRNA contour length (5.4 μm), or from a possible polysome hairpin, which would be approximately 2.7 μm. We estimate the degree of compaction for the AHNAK mRNA relative to its contour length is about 27-fold (using the median distance between the 5’ and 3’ ends compared to the extended contour length) or 18-fold (using the median distance between one end and the middle relative to half the contour length). We obtained similar results for the DYNC1H1 mRNA with the median compaction relative to DYNC1H1 mRNA contour length estimated to be between 21- or 12-fold (Supplemental Figure 4B, C). Similar compaction values were also observed for three other long mRNAs in translating conditions by Srivathsan et al., 2018. These results show that at least these long mRNAs are not in an extended conformation even when engaged in translation and suggest possible mechanisms of mRNA compaction (see discussion).

### AHNAK and DYNC1H1 mRNPs compact further under stress conditions

A similar analysis during stress conditions revealed that the distances between all three smFISH spots for AHNAK and DYNC1H1 mRNAs shrink considerably under arsenite-treated conditions (Figure 2D, E, Supplemental Figure 4D, E). For example, the median distance between the 5’ and 3’ ends, 5’ end and middle, and 3’ end and middle of AHNAK mRNAs are now ~ 80 nm, ~110 nm, and ~ 90 nm respectively. Relative to the contour length, the median compaction of AHNAK mRNAs in U-2 OS cells treated with arsenite is ~ 67.5- or 27-fold; ~67.5-fold if we measure the compaction by dividing the median end-to-end distances to the contour length and ~27-fold if we measure the compaction by dividing the median end-to-middle distances to half the contour length. Similar compaction values were also seen with DYNC1H1 mRNAs (Supplemental Figure 4D, E). The distances between all three smFISH spots for AHNAK and DYNC1H1 mRNAs also shrink considerably in heat shock conditions compared to non-stressed U-2 OS cells, with similar compaction values (Figure 2F, G, Supplemental Figure 4F, G). Thus, in multiple stresses that inhibit translation initiation, we and others (Srivathsan et al., 2018) observe enhanced mRNP compaction.

### Increased compaction of AHNAK and DYNC1H1 mRNPs under stress is a consequence of translational inhibition

Since 80% of AHNAK and 53% of DYNC1H1 mRNAs are found in arsenite-induced SGs at 60’ and similar numbers were seen for heat shock-induced SGs at 60’ (Khong et al., 2017), we expect the compaction measurements for AHNAK and DYNC1H1 mRNAs are reflective of AHNAK and DYNC1H1 mRNAs found inside SGs. Due to technical limitations, we have not been able to examine all three smFISH probes simultaneously with an SG marker. However, smFISH staining indicates most AHNAK and DYNC1H1 mRNAs tend to cluster during stress as expected by their strong enrichment in SG (Figure 2D, F, Supplemental Figure 4D, F). This suggests most AHNAK and DYNC1H1 mRNPs form compact assemblies inside SGs during a stress response.

In principle, the increased compaction of AHNAK and DYNC1H1 mRNPs in SGs might be a consequence of translation inhibition and/or a consequence of increased macromolecular crowding possibly occurring inside SGs compared to the cytosol. We performed two analyses to distinguish these possibilities. First, we measured the distances between the 5’ and 3’ ends of AHNAK and DYNC1H1 mRNPs inside and outside SGs (Figure 3E). We observed that the distances between the 5’ and 3’ ends of AHNAK and DYNC1H1 mRNPs showed a similar distribution inside and outside SGs (Figure 3E), consistent with the increased compaction being independent of SG accumulation.

**Figure 3.**
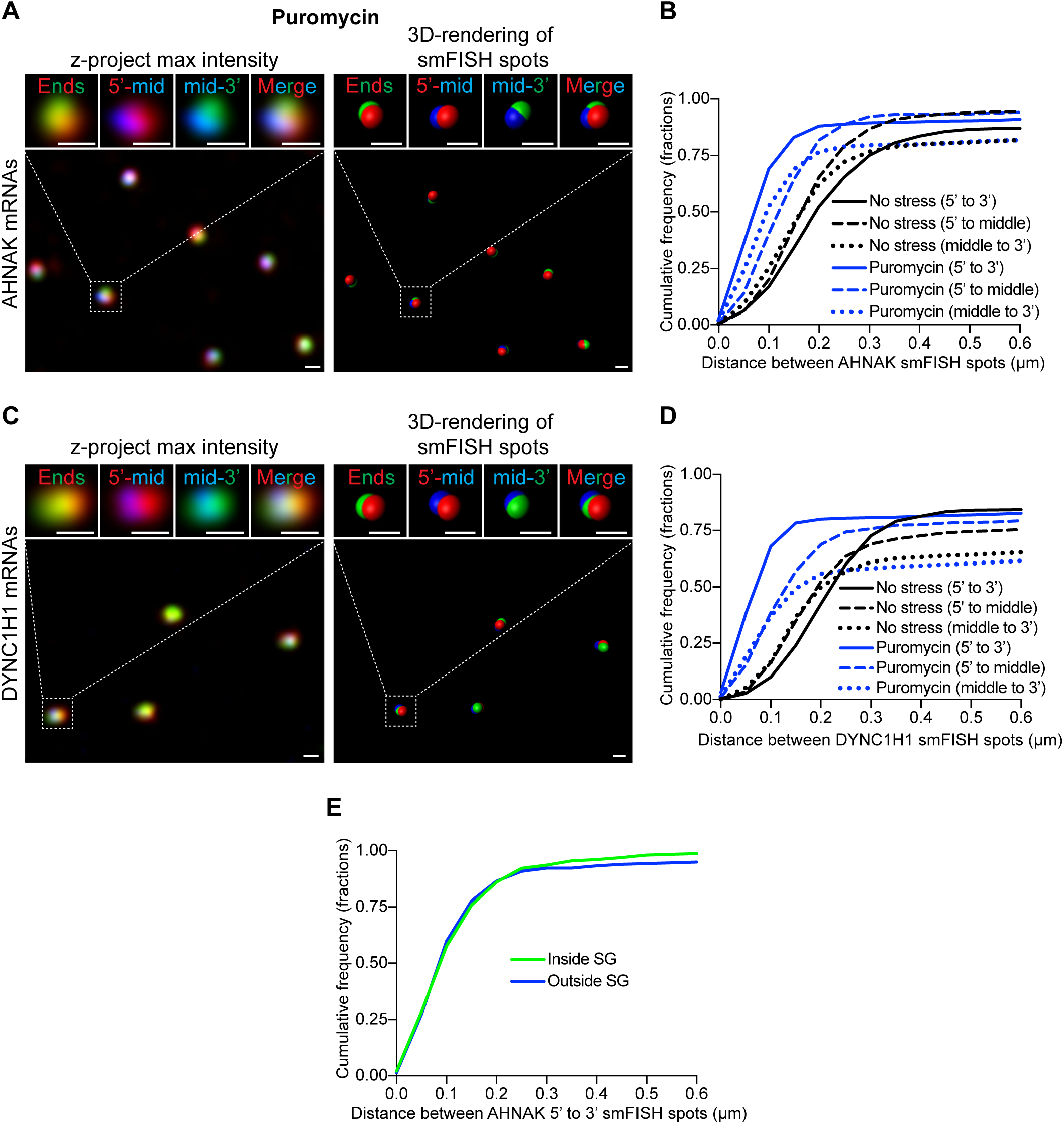
AHNAK and DYNC1H1 mRNPs compact when U-2 OS cells are treated with puromycin. **(A, C)** Left panels. Representative AHNAK and DYNC1H1 smFISH images of U-2 OS cells treated with 10 μg/mL puromycin for one hour. Cells were stained with smFISH probes that bind specifically to the 5’ end (false-colored red), middle (false-colored blue), and 3’ end (false-colored green) of AHNAK and DYNC1H1 mRNAs. Right panels. 3D rendering of smFISH spots by Bitplane Imaris imaging analysis software. Scale bar: 250 nm. **(B, D)** Cumulative frequency graphs (in fractions) of smallest distances between 5’ to 3’ end smFISH spots (solid lines), 5’ end to middle smFISH spots (dash lines), and middle to 3’end smFISH spots (dotted lines) in unstressed cells (black) and 10 μg/mL puromycin-treated cells (blue). More than 1100 smallest distances were quantified for each sample. **(E)** Cumulative frequency graph (in fractions) of smallest distances between 5’ to 3’ end smFISH spots (solid lines) inside and outside SG in U-2 OS cells stressed with 60’ 0.500 mM NaAsO_2_. More than 500 smallest distances were quantified. The analysis was performed with the experimental results as shown in Figure 2B-C.

In a second experiment, we examined AHNAK and DYNC1H1 mRNPs organization by smFISH when U-2 OS cells were treated with puromycin (Figure 3A-D). Puromycin leads to release of elongating ribosomes but does not lead to SG assembly, perhaps because translation initiation is ongoing and even partial ribosome engagement appears to block mRNA accumulation in SG (see below). We observed puromycin is sufficient to lead to increased compaction of the AHNAK and DYNC1H1 mRNPs (Figure 3B, D), even though SG do not form. Similar compaction of the MDN1, POLA1, and PRPF8 mRNPs with puromycin treatment have been reported in HEK293T cells (Srivathsan et al., 2018). These results argue that mRNP compaction is not due to increased macromolecular crowding found inside SGs and instead is a consequence of translational inhibition and the loss of elongating ribosomes.

### Compaction of mRNPs during stress proceeds in a 5’ to 3’ direction

Since mRNP compaction is likely a consequence of translation inhibition, we hypothesized that the compaction of AHNAK and DYNC1H1 mRNPs under stress is mediated by intramolecular interactions formed within the ORF of mRNAs in the absence of ribosomes. If this model is accurate, mRNP compaction will begin at the 5’end of the transcript as elongating ribosomes translocate towards the 3’end of the transcript once translation initiation is inhibited. This model predicts that the 5’end to the middle of the mRNA will compact first as elongating ribosomes move down the mRNA in the absence of new translation initiation, followed by a subsequent compaction of the middle and the 3’end of AHNAK mRNPs as ribosomes finally exit the ORF and terminate translation. To examine this possibility, we stressed U-2 OS cells for 10, 20, and 30 minutes with 500 μM arsenite and examined the distances between the different regions of the AHNAK mRNA by smFISH (Figure 4).

**Figure 4.**
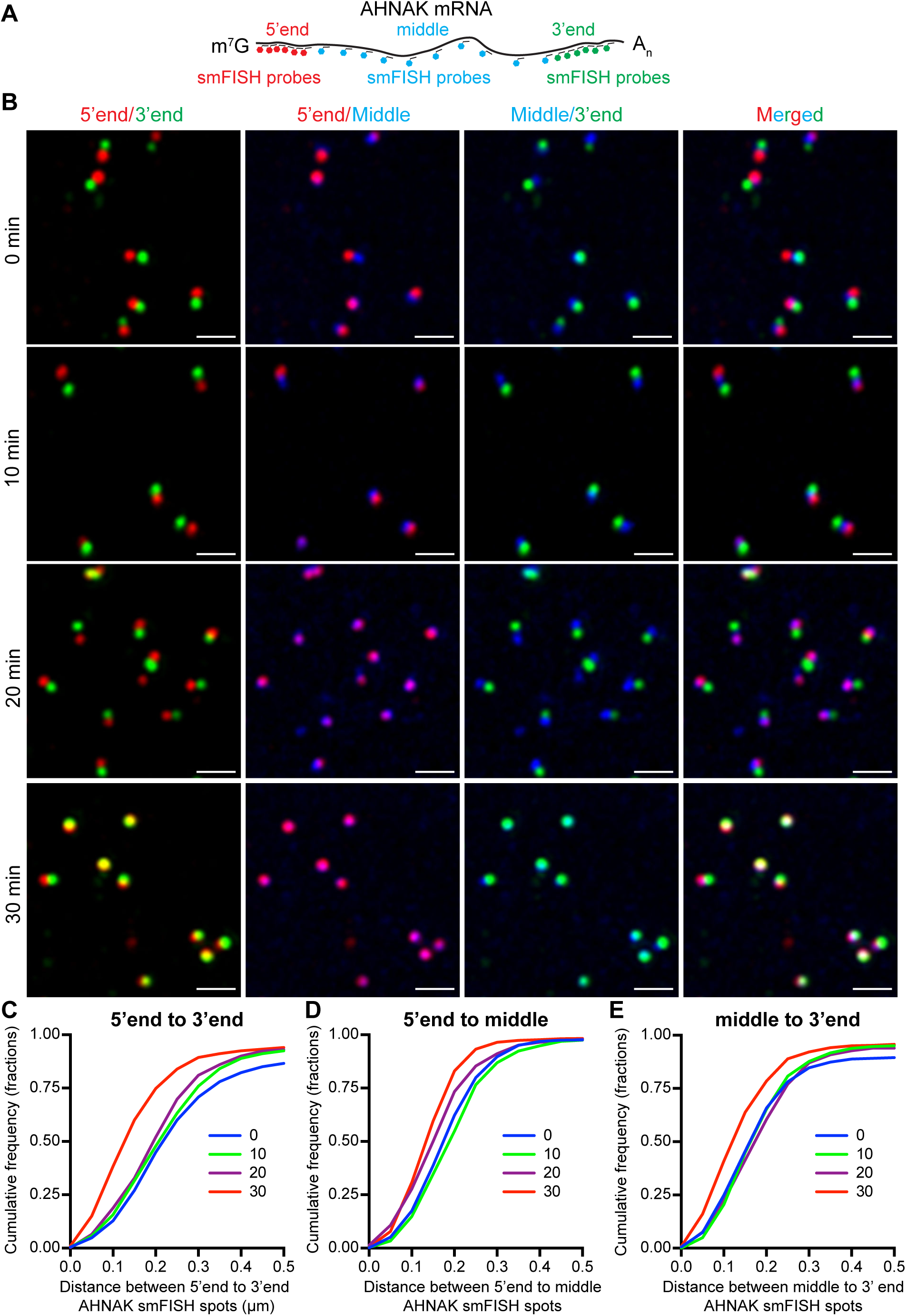
Distances between the 5’end and middle AHNAK smFISH spots shrink first after the addition of NaAsO_2_. **(A)** Cartoon schematic illustrating where smFISH probes bind to AHNAK mRNAs. smFISH probes binding to the 5’end, middle or 3’end are labeled with distinct fluorophores and are false-colored as red, blue, and green respectively. **(B)** Representative AHNAK smFISH image of U-2 OS cells that were not stressed or stressed with 0.5 mM NaAsO_2_ for 10, 20 and 30 minutes. Scale bar: 1 μm. **(C-E)** Cumulative frequency graphs (in fractions) of smallest distances between **(C)** 5’ to 3’ end smFISH spots, **(D)** 5’ end to middle smFISH spots and **(E)** middle to 3’end smFISH spots in unstressed U-2 OS cells or 0.5 mM NaAsO_2_-treated U-2 OS cells for 5-30 minutes. More than 800 smallest distances were quantified for each sample.

Qualitatively, we observed the distance between 5’end and the middle are closer at 20 minutes after addition of arsenite, at which time the distance between the 5’end and 3’end or the middle and the 3’end are still separated (Figure 4B). Quantitatively, the distances between the 5’end and the middle shrink considerably at 20 minutes (Figure 4D), which correlates with the time ribosomes should be beginning to exit the 5’ portion of the coding region since it takes 8 minutes after addition of arsenite to maximize eIF2a phosphorylation in U-2 OS cells (Wheeler et al., 2016). In contrast, the shrinkage in distances for the 5’end to the 3’end or the middle to the 3’end is only noticeable at 30 minutes (Figure 4C, E). These observations are consistent with the model that intramolecular folding of the mRNA, either through RNA-RNA interactions or protein binding, as ribosomes expose the coding region, leads to the increased compaction of the non-translating mRNA.

### The 5’ and 3’ ends of the AHNAK and DYNC1H1 mRNPs become closer when non-translating

Under non-stress conditions, we notice the 5’ to 3’ end distances of AHNAK and DYNC1H1 mRNPs are larger than one would expect base on specific models of mRNP organization such as the closed-loop model of translation (median ~200 nm) (see discussion). This suggests that the closed-loop model of translation either does not occur on these mRNAs or is transient. However, we noticed that the 5’ and 3’ ends of the AHNAK mRNP and DYNC1H1 mRNPs shrink disproportionally under non-translating conditions and reach a median distance between the ends of ~50 nm (Figure 2E, G, Figure 3B, D, Supplemental Figure 4E, G) These results suggest stress triggers a reorganization of mRNPs that disproportionally brings the 5’ and 3’ ends closer together.

To further examine the relationship between the 5’ and 3’ ends of these mRNAs, we measured the angles between the middle smFISH spots to the 5’ and 3’ ends for AHNAK and DYNC1H1 smFISH spots (Figure 5A). We observed in non-stress conditions, the angles can vary considerably, but most angles are less than 90 degrees with a median angle of ~ 60 degrees (Figure 5B, C). This observation suggests that these mRNAs are not linear in cells and, on average, the 5’ end is closer to the 3’ end than expected by chance, perhaps due to features of polysomes or RNA binding proteins (see discussion).

**Figure 5.**
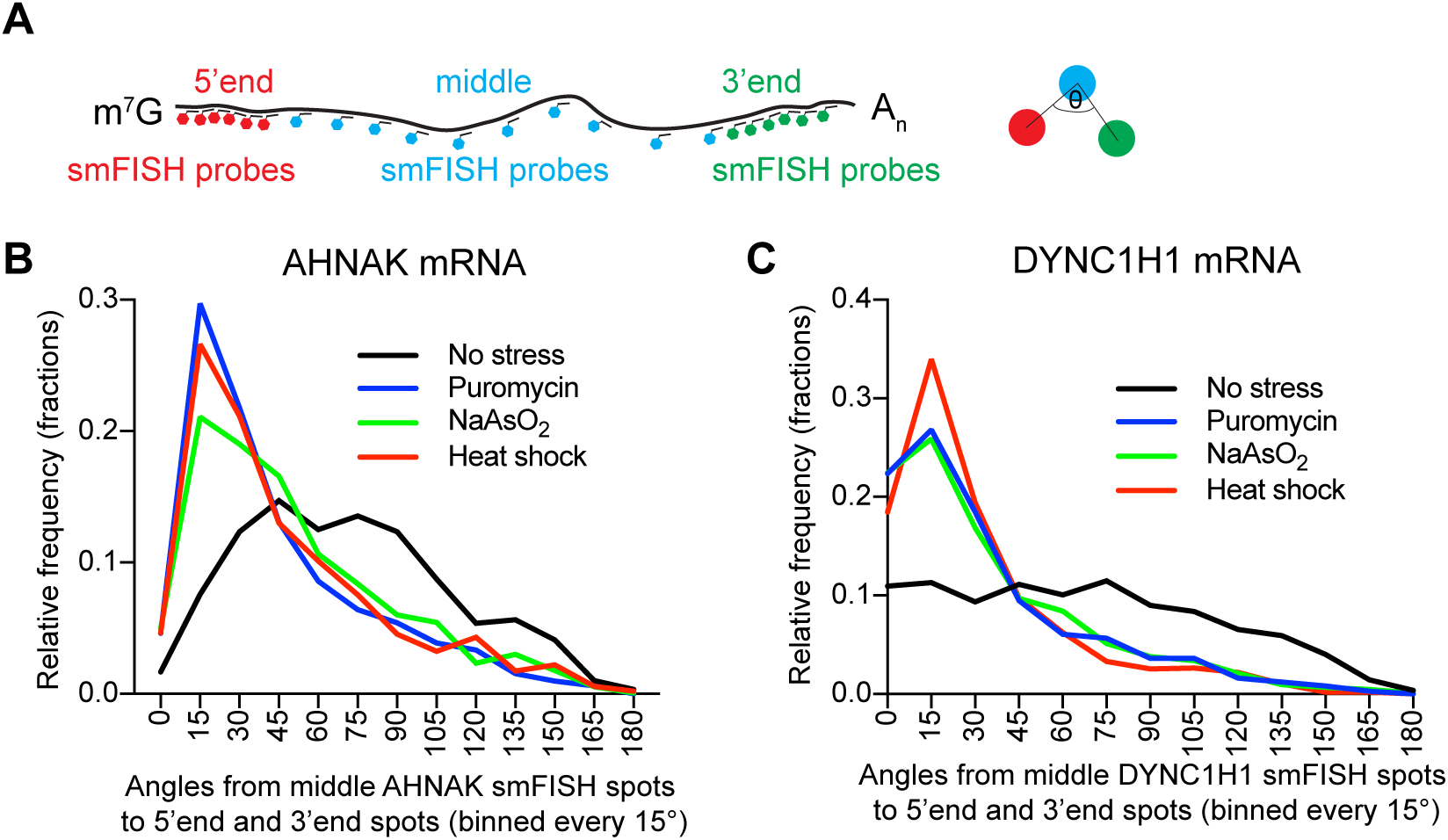
Translation inhibition with puromycin, NaAsO_2_ or heat shock in U-2 OS cells disproportionally shrink the distances between the 5’ and 3’ ends relative the middle of AHNAK and DYNC1H1 mRNPs. **(A)** Cartoon schematic indicating the angles that were measured in **(B, C)**. Histograms illustrating the relative frequency (fractions) of angles from middle smFISH spots to 5’end and 3’end smFISH spots of **(B)** AHNAK and **(C)** DYNC1H1 mRNAs in unstressed (black line), puromycin-treated (blue), NaAsO_2_-treated (green), or heat shocked (red) U-2 OS cells. The histograms were generated by binning every 15°. More than 850 angles were quantified for each sample.

A striking result was that under non-translating conditions, most angles are now less than 45 degrees with a median angle of 20 degrees (Figure 5B, C). Since the mRNAs are now compact under these conditions, we were concerned that the small spacing between the smFISH spots might skew this analysis. Given this, we performed a second analysis where we limited our analysis to specific mRNAs where the total distance between the 5’end to middle and the middle to 3’end is between 0.3 μm to 0.6 μm with the goal of increasing our ability get an accurate angle measurement. This analysis also showed a dramatic reduction in the angle between the 5’-middle-3’ signals (Supplemental Figure 5). This provides a second line of evidence that when mRNAs exit translation the distance between the 5’ and 3’ ends of AHNAK and DYNC1H1 mRNPs is notably compacted, perhaps due to the polymer nature of the mRNA (see discussion).

## DISCUSSION

### mRNAs need to exit translation completely before entering SGs

We present several observations that indicate mRNAs must be completely disengaged from translating ribosomes before entering SGs (Figure 6). First, mRNAs with long ORF are slower at recruitment to SGs than mRNAs with short ORF (Figure 1A, B, Supplemental Figure 1). Second, recruitment of mRNAs with long ORF to arsenite-induced SGs is quicker when puromycin is added, which will rapidly disengage elongating ribosomes (Figure 1A, C, Supplemental Figure 2). Third, addition of cycloheximide to cells treated with arsenite for 30 minutes inhibits additional recruitment of RNA to SGs (Figure 1A, D, Supplemental Figure 3), which indicates that the continued accumulation of AHNAK and DYNH1C1 mRNAs require ribosome run-off. For mRNAs with long ORF such as AHNAK and DYNH1C1, a large amount of the ORF will be exposed by 30 minutes of stress, yet these mRNAs have only partially accumulated in SG. This argues that mRNAs must fully disengage from elongating ribosomes before stable accumulation in SGs. Additional evidence in support of this model comes from single-molecule experiments that show mRNAs engaged with ribosomes can only form a transient association with SG and do not enter a stable association, which can be seen with non-translating mRNAs (Moon et al., 2018).

**Figure 6.**
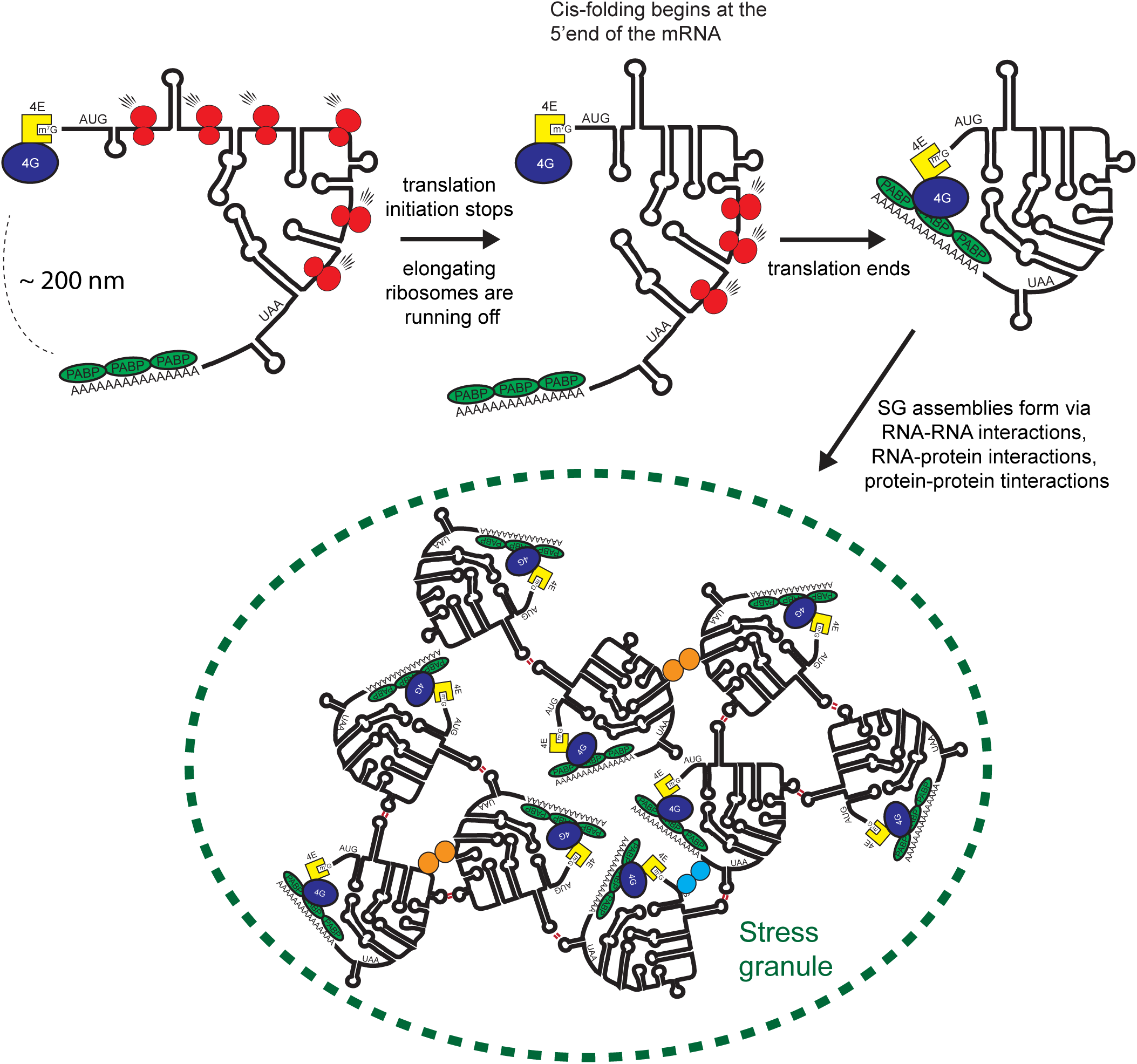
Model depicting mRNP compaction and mRNA recruitment to SGs. Under non-stress conditions, mRNPs are engaged in translation. Relative to its contour length, significant compaction was observed. During early stages of stress, ribosomes will migrate towards the 3’end of mRNAs and mRNAs start to compact at the 5’end, most likely due to intramolecular interactions formed. When mRNA exits translation, the mRNP compacts further and the ends are disproportionally close. One hypothesis is the closed-loop conformation is re-established. Some of these mRNPs, preferentially long mRNAs, start to accumulate in SGs via intermolecular interactions formed with other mRNPs.

It is an unsolved mystery why complete ribosome disengagement is required for stable association of mRNAs with SG. One possibility is that the mRNA association with the translation machinery increases the presence of helicases and/or protein chaperones that prevent or disengage interactions between the translating mRNP and SG. Alternatively, it may be energetically unfavorable for an 80S ribosome and its associated factors to enter the altered environment of an SG, either because of energetic costs of changes in solvation, or because the mesh size of an SG is smaller than an assembled 80S ribosome.

### mRNPs are compact in both translating and non-translating states

We present several observations that demonstrate mRNAs are compacted >10-fold relative to its contour length even when it is translating. First, under no stress conditions, the median distances between the 5’ and 3’end of AHNAK and DYNC1H1 mRNPs are roughly ~150 nm (Figure 2C, Supplemental Figure 4C). Relative to the AHNAK and DYNC1H1 mRNAs contour length, 5.6 and 4.2 μm respectively, this is ~27- and ~21-fold compaction respectively. Second, the median distances between the 5’ and mid or 3’ and mid of AHNAK and DYNC1H1 are roughly ~100 nm (Figure 2C, Supplemental Figure 4C). Relative to half of its contour length, this is ~18- and ~12-fold compaction. Third, we notice the angles between the middle and the 5’ and 3’end of AHNAK and DYNC1H1 mRNPs are usually less than 90 degrees (Figure 5B, C, Supplemental Figure 5B, C) which also suggests significant compaction of the ends relative to its linear length. Similar results are seen for the MDN1, POLA1, and PRPF8 in HEK293T mRNAs in HEK293 cells indicating this is a general phenomenon (Srivathsan et al., 2018).

We suggest two mechanisms account for the compaction of mRNAs during non-stress conditions. First, we suggest transient folding of the ORF region between elongating ribosomes compacts mRNAs (Figure 6). The average inter-ribosome distance is estimated to be 150 nucleotides in yeast and 189 in mammalian cells (Arava et al., 2003; Hendrickson et al., 2009), or from single-molecule translation assays in mammalian cells, the average inter-ribosomal is estimated to be between 200-900 nucleotides (Morisaki et al., 2016; Wang et al., 2016; Wu et al., 2016; Yan et al., 2016). Additionally, for DYNC1H1 mRNPs specifically, it is estimated that on average each DYNC1H1 mRNA has about 7 ribosomes (Pichon et al., 2016). Since a ribosome footprint is ~30 nucleotides (Steitz, 1969; Wolin and Walter, 1988), this suggests most of the nucleotides in the ORF are not covered with ribosomes. We estimate ~80-97% of the ORF nucleotides for most mRNAs, and ~98% for DYNC1H1 mRNA, are not engaged with ribosomes. Therefore, the ORF region can form significant intramolecular interactions with itself, or with the 5’ and 3’ UTRs. Supporting this model, an extensive physical association between the 3’UTR and the ORF has been reported for mRNAs (Eldad et al., 2008). Besides intramolecular interactions, the folding of the ORF region may also be promoted by RNA-binding proteins or complexes by connecting different ORF regions of the mRNA.

A second mechanism of compacting translating mRNAs may arise from the architecture of polysomes since the path a mRNA takes within each ribosome is curved (Agrawal et al., 1996). Therefore, by its nature, a translating mRNA would be more compact compared to its contour length. Indeed, instances of circular, spiral, rosette, staggered line, double-row and helical polysomes has been observed by traditional EM or more advanced cryo-EM and cryo-ET methods, all of which would compact the overall shape of the mRNA (Afonina et al., 2014; Afonina et al., 2015; Afonina Zh et al., 2013; Brandt et al., 2010; Brandt et al., 2009; Daneholt et al., 1977; Kopeina et al., 2008; Madin et al., 2004; Myasnikov et al., 2014; Palade, 1955; Viero et al., 2015; Warner et al., 1962; Wettstein et al., 1963; Yazaki et al., 2000).

We, and others (Adivarahan et al., 2017), observe under stress conditions when mRNAs stop translating, mRNPs compact further (Figure 6). Specifically, the distances between the 5’ to 3’ends, 5’end to the middle and 3’end to the middle are smaller for AHNAK and DYNC1H1 mRNAs under a variety of conditions that cause mRNAs to disengage from elongating ribosomes, such as arsenite, heat-shock, and puromycin (Figure 2, Figure 3, Supplemental Figure 4). This additional compaction appears a consequence of translational shutoff and not a consequence of being inside SGs for two reasons (Figure 6). (1) Compaction is similar inside and outside SGs (Figure 3E). And (2) puromycin also compacts AHNAK and DYNC1H1 mRNPs without inducing SG (Figure 3A-D). The most straightforward interpretation for increased mRNP compaction during stress is mRNAs forming increased intra-molecular interactions in the absence of translating ribosomes. This interpretation is supported by the fact that the compaction precedes temporally in a 5’ to 3’ manner that correlates with ribosomes transiting towards the 3’end of AHNAK mRNAs after addition of arsenite (Figure 4).

### The spatial relationship between the 5’ and 3’ ends change with stress

We observed mRNPs are reorganized during stress in a manner where the distances between the ends are now smaller than the distances between the ends to the middle for AHNAK and DYNC1H1 mRNAs (Figure 6). Specifically, the median distance between the 5’ and 3’end is ~ 50 nm during stress while the median distance between the 5’ end to the middle or 3’ end to the middle is ~ 100 nm (Figure 2E, G, Figure 3B, D, Supplemental Figure 4E, G). This is different with respect to translating mRNPs; the median distance between the ends (~ 200 nm) is larger than the median distance between the 5’end or 3’end to the middle (~150 nm) (Figure 2C, Supplemental Figure 4C). In support of the 5’ and 3’ ends being in proximity under stress, we also observed the angles between the middle and the ends of AHNAK and DYNC1H1 mRNAs are now considerably smaller under stress (Figure 5B, C, Supplemental Figure 5B, C).

We suggest two possible mechanisms for why the 5’ and 3’ ends may enter into proximity during stress. One hypothesis is that the closed-loop conformation is a non-stable state during translation and that in the absence of translation, the closed loop confirmation can form through interactions of eIF4E, eIF4G, and PABP where eIF4E binds to the m^7^G cap, PABP binds to poly(A) tail, and eIF4G binds to both eIF4E and PABP (Hinnebusch and Lorsch, 2012) (Figure 6). Alternatively, or in addition, the 5’ and 3’ ends of mRNPs may be close during stress because of an intrinsic property of “naked” RNAs to fold in a manner that brings the ends in proximity. Several computation studies suggest the ends of mRNAs are close (<10 nm) for RNAs in solution (Clote et al., 2012; Fang et al., 2011; Yoffe et al., 2011). Moreover, an *in vitro* FRET-based assay indicates for all eleven mRNAs examined, the distance between the 5’ and 3’ ends are less than 10 nm (Lai et al., 2018). This distance is significantly smaller than one would expect if it these RNAs were behaving as a random coil in solution (Lai et al., 2018). Therefore, under stress conditions, if most mRNAs are now exposed and can form significant intramolecular interactions, its properties as a polymer might promote the interaction of the 5’ and 3’ ends.

To consider whether there could be a direct interaction between the 5’ and 3’ ends of mRNAs at the distances estimated from our smFISH analysis, we estimated what distance we would observe by smFISH for a classic eIF4E-eIF4G-PABP closed loop structure (Supplemental Figure 6). In a closed-loop model with PABP interacting with eIF4G, the distance between the m^7^G-cap and the poly(A) tail ends should be less than 20 nm since the diameter of an average protein is about ~5 nm (Milo et al., 2010). We estimate the distance between the m^7^G-cap and the last nucleotide that precedes the poly(A) tail should be ~50 nm, since the average poly(A)-tail of a mammalian mRNA is < 100 nucleotides (Chang et al., 2014), and when fully extended is ~30 nm in length (Milo et al., 2010). Finally, given where the 5’ and 3’end smFISH probes bind on AHNAK and DYNC1H1 mRNAs and provided if the overall compaction of AHNAK and DYNC1H1 mRNAs is similar at the ends (>20 fold), we estimate a distance less than 80-65 nm between 5’end and 3’end of AHNAK and DYNC1H1 smFISH spots respectively could support a closed-loop conformation (Supplemental Figure 6). Although these calculations should be taken as ballpark estimates, this would suggest that less than 20% of AHNAK and DYNC1H1 translating mRNPs have distances between the ends supporting a closed-loop conformation. In contrast, during stress, when mRNAs exit translation, >50% of AHNAK and DYNC1H1 mRNAs have distances that are consistent with the closed loop-conformation (Figure 2C, E, G, Figure 3B, D, Supplemental Figure 4C, E, G). Similar results have also been described for the MDN1, POLA1, and PRPF8 mRNAs (Adivarahan et al., 2017) suggesting this effect is not limited to the mRNAs we have examined. Thus, one mechanism for the shortened distance between the 5’ and 3’ ends during stress could be direct protein-protein interactions (Figure 6).

The observation that the distances between the ends of translating mRNPs are typically large (greater than 100 nm) is surprising with respect to many aspects of established RNA biology. For example, the closed loop model as discussed earlier but also for other 3’UTR regulatory elements that can affect processes occurring at the 5’UTR (e.g. miRNA-mediated translation initiation repression). Our observations suggest that this is not physically possible for translating mRNPs unless there is a large network of protein-protein interactions that connect the ends (>20 proteins since an average protein size is 5 nm). Alternatively, we hypothesize, based on observations derived from non-translating conditions, effects imparted by 3’UTR regulatory elements on processes at the 5’end will only occur when all translating ribosomes are released from the mRNA. If this is accurate, this suggests mRNAs that are being translated are likely unaffected by these 3’UTR regulatory elements. However, when mRNA loses all its translating ribosomes, most likely in a stochastic manner, these regulatory elements can now communicate with the 5’end and affect mRNA fate. This leads to a model wherein 3’ UTR elements can affect events at the 3’ end of the mRNA, such as deadenylation, regardless of translation status, but the ability of the 3’ UTR and poly(A) tail to influence events at the 5’ mRNA end, such as translation initiation and decapping, would be more pronounced on mRNAs that have exited translation.

## Materials and Methods

### U-2 OS growth conditions

Human osteosarcoma U-2 OS cells (Kedersha et al., 2016), maintained in DMEM with high glucose, 10% fetal bovine serum and 1% penicillin/streptomycin at 37°C/5% CO_2_, were used in all experiments.

### Sequential immunofluorescence and single molecule FISH

The protocol was performed as described previously (Khong et al., 2018; Khong et al., 2017). Briefly, U-2 OS cells were seeded on sterilized coverslips in 6-well tissue culture plates. At ~80% confluency, media was exchanged sixty minutes before experimentation with fresh media. Experimentation was performed as described in each figure. U-2 OS cells were treated with 500 μΜ NaAsO2, 10 μg/mL puromycin or 50 μg/mL cycloheximide as described. After stressing cells, the media was aspirated and the cells were washed with pre-warmed 1x PBS. The cells were then fixed with 500 μL 4% paraformaldehyde for ten minutes at room temperature.

After fixation, cells were washed twice with 1x PBS, permeabilized in 0.1% Triton X-100 1x PBS for five minutes, and washed once with 1x PBS. Coverslips were transferred to a humidifying chamber and cells were incubated in 5 μg/mL mouse α-G3BP1 antibody (ab56574, Abcam) in 1x PBS for sixty minutes at room temperature. Afterward, the coverslips were transferred to a 6-well plate and washed three times with 1x PBS. Coverslips were then transferred back to the humidifying chamber and incubated in goat α-mouse FITC-conjugated antibody in 1× PBS (1:1000 dilution ab6785, Abcam) for sixty minutes at room temperature. The coverslips were transferred to 6-well plate and washed three times with 1x PBS. Antibodies binding to cells were fixed on cells by incubating coverslips with 500 μL 4% paraformaldehyde for ten minutes at room temperature.

After immunofluorescence, smFISH was performed as described previously (Khong et al., 2018) using Biosearch Technologies Stellaris buffers (SMF-HB1-10, SMF-WA1-60, SMF-WB1-20). Specific smFISH probes were created using software designed by Biosearch Technologies (https://www.biosearchtech.com/support/tools/design-software/stellaris-probe-designer). smFISH probes that bind to AHNAK, DYNC1H1, NORAD, PEG3, ZNF704, and CDK6 mRNAs were designed previously (Khong et al., 2017; Moon et al., 2018). Newly designed smFISH probes include probes that bind to the middle of AHNAK mRNAs, and 5’end, middle and 3’end of DYNC1H1 mRNAs. The smFISH probes were made by conjugating 30 or 60 DNA-oligos with ddUTP-Atto488, ddUTP-Atto550, or ddUTP-Atto633 as described in Gaspar et al., (2017).

### Imaging parameters

Fixed stained U-2 OS cells were imaged using a wide-field DeltaVision Elite microscope with a 100x objective using a PCO Edge sCMOS camera with appropriate filters as described previously (Khong et al., 2018). At least thirty Z-sections (0.2 μm step size) were captured for each image in order to capture the entire U-2 OS cell. Imaging parameters were adjusted to capture fluorescence within scope’s dynamic range. After images were collected, the images were then deconvolved with built-in DeltaVision software as described previously (Khong et al., 2018).

### Image analysis

All image analysis was performed using Bitplane Imaris image analysis software as described previously with the deconvolved images (Khong et al., 2018). To measure the fraction of smFISH spots in SGs in U-2 OS cells, please refer to Khong et al., 2018.

We quantified the distances between the 5’end, middle, and 3’ end of AHNAK and DYNC1H1 mRNAs with the help of Bitplane Imaris Imaging Analysis software in the following manner. (1) First, we open the deconvolved DeltaVision images in Bitplane Imaris Imaging Analysis Software (see imaging parameters). Bitplane Imaris Imaging Analysis Software reassembles the Z-stack DeltaVision images in 3-D automatically. (2) Second, we mask all fluorescent signal coming from the nuclei of cells of an image using DAPI staining to define the nuclei. (3) Third, we applied the spot creation wizards to identify the 5’end, middle, and 3’end AHNAK and DYNC1H1 smFISH spots using these two following parameters: a fixed xy diameter spot size of 200 nm and a manually determined fluorescent quality threshold. Upon identification of smFISH spots, the spot creation wizard provides all x,y,z coordinates for the center of each smFISH spot in an excel spreedsheet. (4) Fourth, we exported the coordinates of all smFISH spots and computed the distances between all smFISH spots in different channels by applying the distance formula between two points in 3D-space. (5) Finally, we note the smallest distance between all smFISH spots of different channels. We assume the smallest distance is the distance between two regions of a single AHNAK or DYNC1H1 mRNA molecule.

With respect to angles, with the smallest distances between smFISH spots provided, we computed the angles between the middle smFISH spots to the 5’ and 3’end smFISH spots by applying the law of cosines with three known sides. We only computed the angles when the smallest distance measured between the three smFISH spots (5’end, middle, and 3’end) can be attributed to all three smFISH spots.

## ACKNOWLEDGMENTS

We would like to thank Nancy Kedersha and Paul Anderson for the U-2 OS cells. We would like to thank Anne Ephrussi for providing ddUTP-Atto633 and Evan Lester for making CDK6 smFISH probes. We would like to thank Carolyn Decker for DeltaVision training. We are grateful to Joe Dragavon for image analysis training (Bitplane Imaris Image Analysis Software). The imaging data analysis was performed at the CU Light Microscopy Core Facility and the BioFrontiers Institute Advanced Light Microscopy Core. We also like to thank Theresa Nahreini for cell culture facility training. All cell culture experiments conducted was at BioFrontiers Cell Culture Facility. Finally, we would like to thank Olke Uhlenbeck for helpful discussion. This work was funded by NIH grants GM045443 (R.P.) and the Howard Hughes Medical Institute (R.P.).

---

**Supplemental Figure 1.**
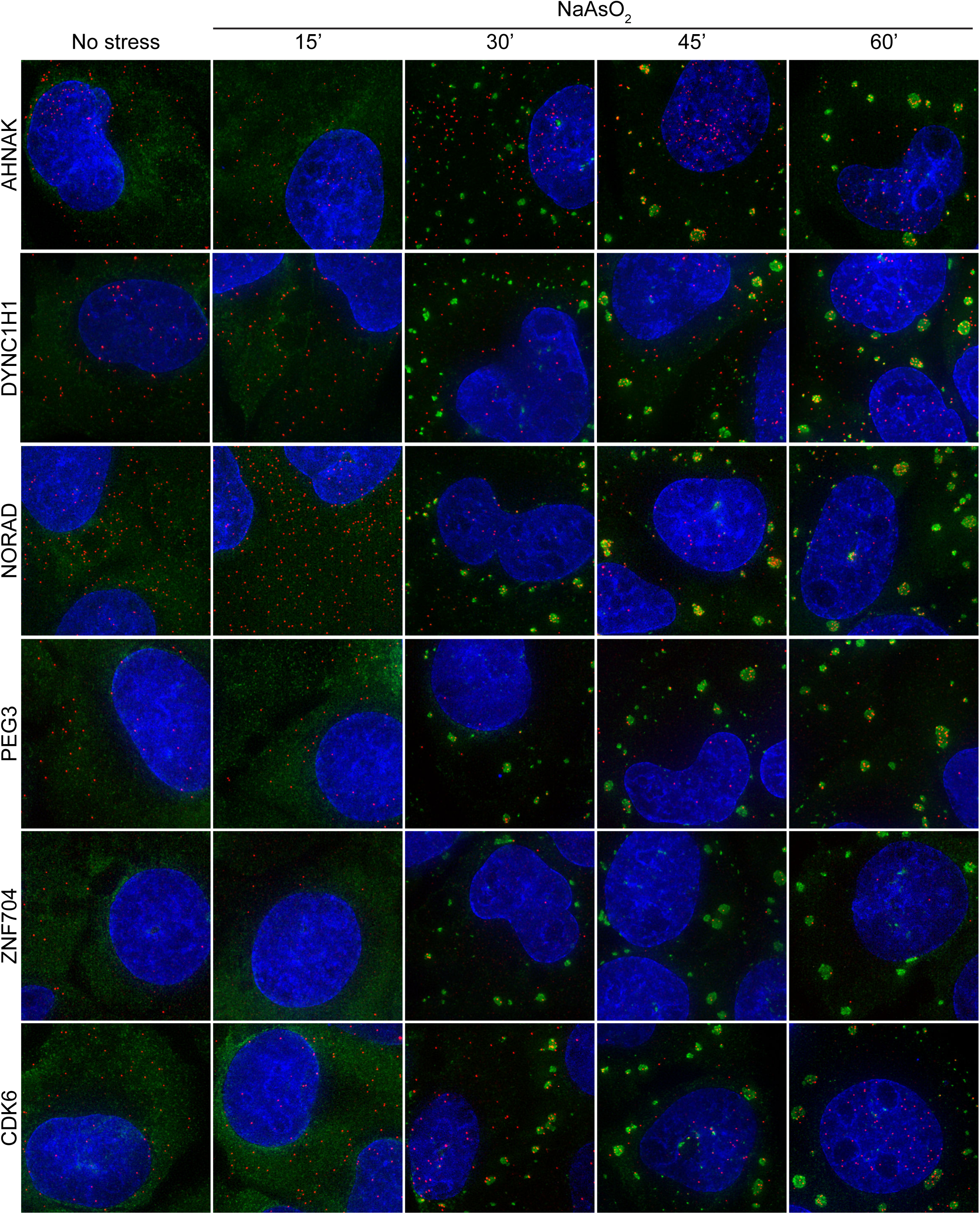
When mRNAs are recruited SGs is correlated with ORF length in U-2 OS cells. Representative smFISH images acquired for six different transcripts (AHNAK, DYNC1H1, NORAD, PEG3, ZNF704, CDK6) for U-2 OS cells treated with 0.5 mM NaAsO_2_ for 15, 30, 45, or 60 minutes. Cells were stained with Nuclei (false-colored blue), G3BP1 (false-colored green), and specific transcripts by smFISH (false-colored red).

**Supplemental Figure 2.**
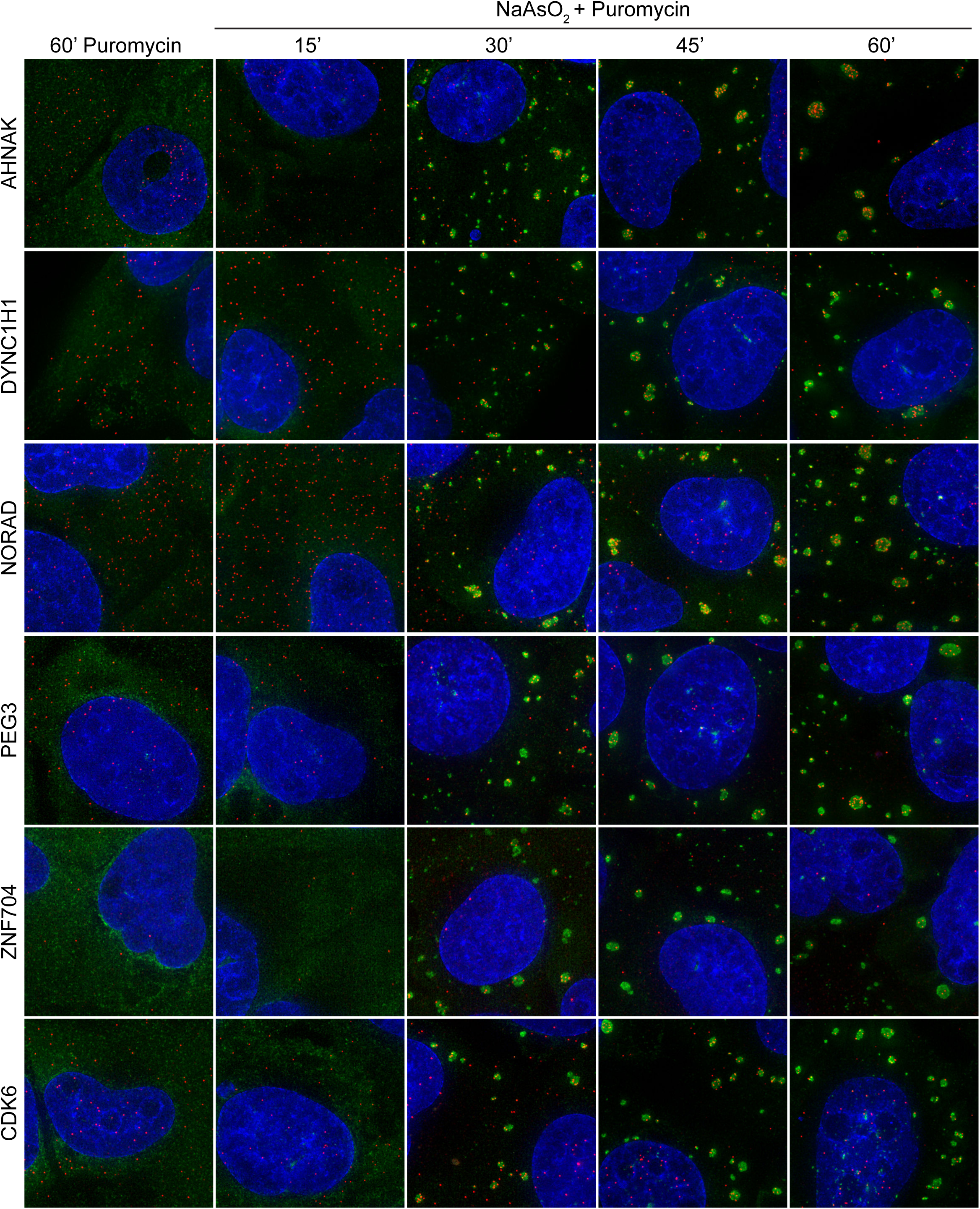
mRNAs with long ORF are recruited to SGs quicker with puromycin in U-2 OS cells. Representative smFISH images acquired for six different transcripts (AHNAK, DYNC1H1, NORAD, PEG3, ZNF704, CDK6) for U-2 OS cells treated with 10 μg/mL puromycin and 0.5 mM NaAsO_2_ for 15, 30, 45, or 60 minutes. Cells were stained with Nuclei (false-colored blue), G3BP1 (false-colored green), and specific transcripts by smFISH (false-colored red).

**Supplemental Figure 3.**
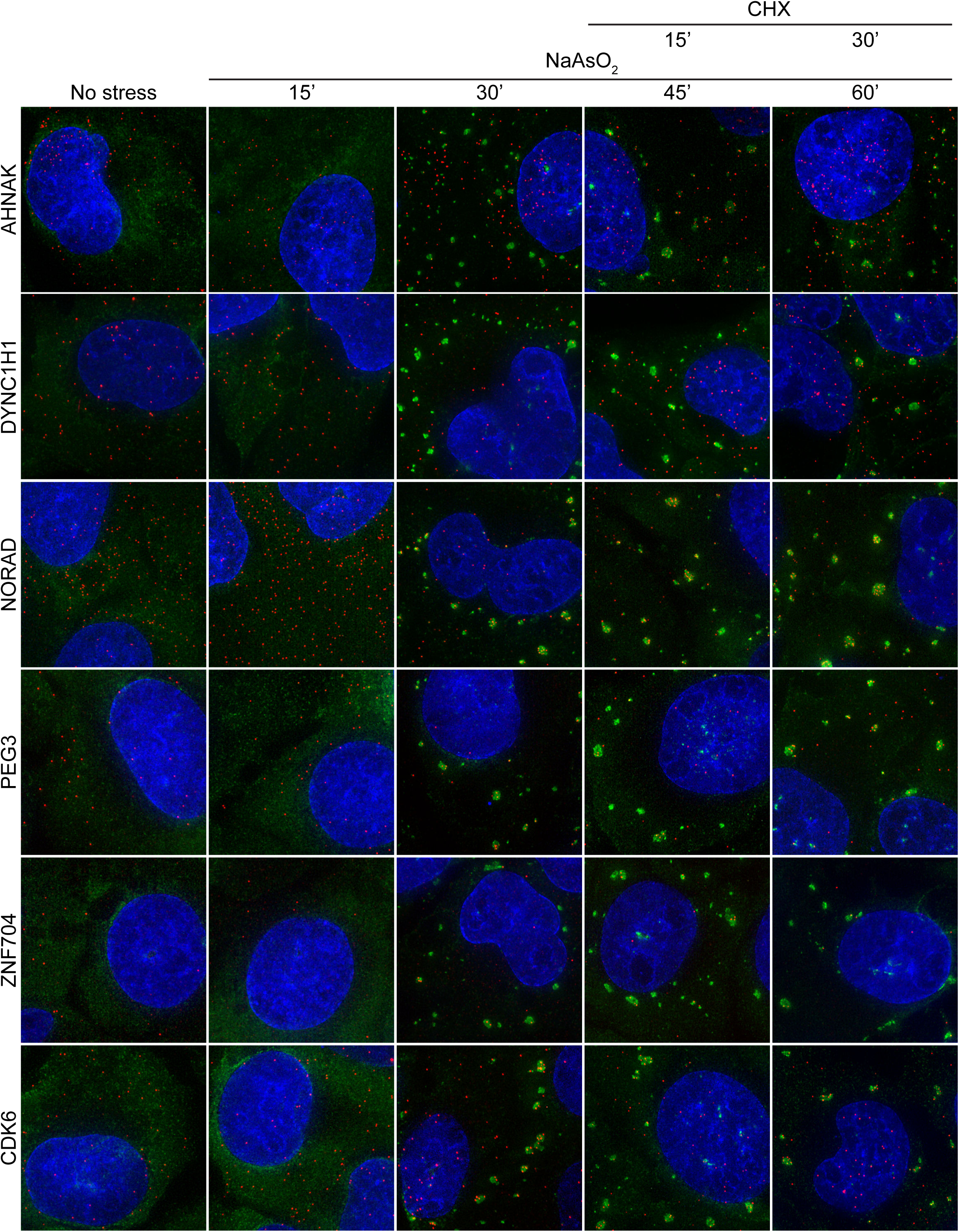
Cycloheximide added at 30 minutes after treating U-2 OS cells with NaAsO_2_ impedes recruitment of mRNA to SGs. Representative smFISH images acquired for six different transcripts (AHNAK, DYNC1H1, NORAD, PEG3, ZNF704, CDK6) for U-2 OS cells treated with 0.5 mM NaAsO_2_ for 15, 30, 45, or 60 minutes with 50 μg/mL cycloheximide added at 30 minutes. Cells were stained with Nuclei (false-colored blue), G3BP1 (false-colored green), and specific transcripts by smFISH (false-colored red).

**Supplemental Figure 4.**
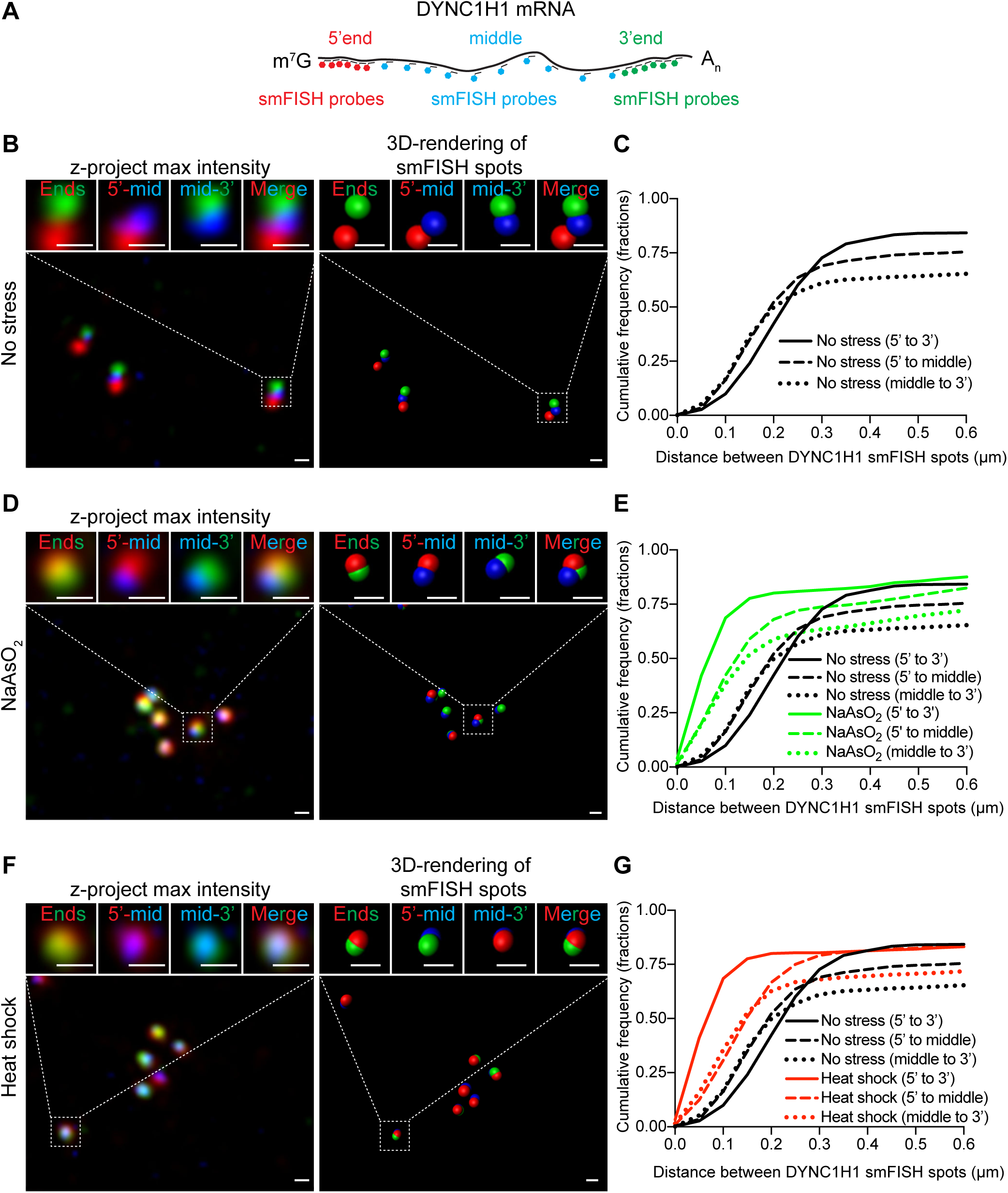
DYNC1H1 mRNPs organization in non-stress and stress conditions. **(A)** Cartoon schematic indicating where smFISH probes bind to DYNC1H1 mRNAs. smFISH probes binding to the 5’ends, middle or 3’ends are labeled with distinct fluorophores and are false-colored red, blue, and green respectively. **(B, D, and F)** Representative DYNC1H1 smFISH images of U-2 OS cells that were **(A)** not stressed or **(C)** stressed with 0.500 mM NaAsO_2_ for 60’ or **(E)** heat shock at 42°C for 60’. Cells were stained with DYNC1H1 smFISH probes that bind specifically to the 5’ end (false-colored red), middle (false-colored blue), and 3’ end (false-colored green). **(B, D, and F)** Cumulative frequency graphs (in fractions) of smallest distances between 5’ to 3’ end smFISH spots (solid lines), 5’ end to middle smFISH spots (dash lines), and middle to 3’end smFISH spots (dotted lines) in unstressed cells (black), 0.500 mM NaAsO_2_-treated cells (green), and heat shock cells (red). More than 1000 smallest distances were quantified for each sample.

**Supplemental Figure 5.**
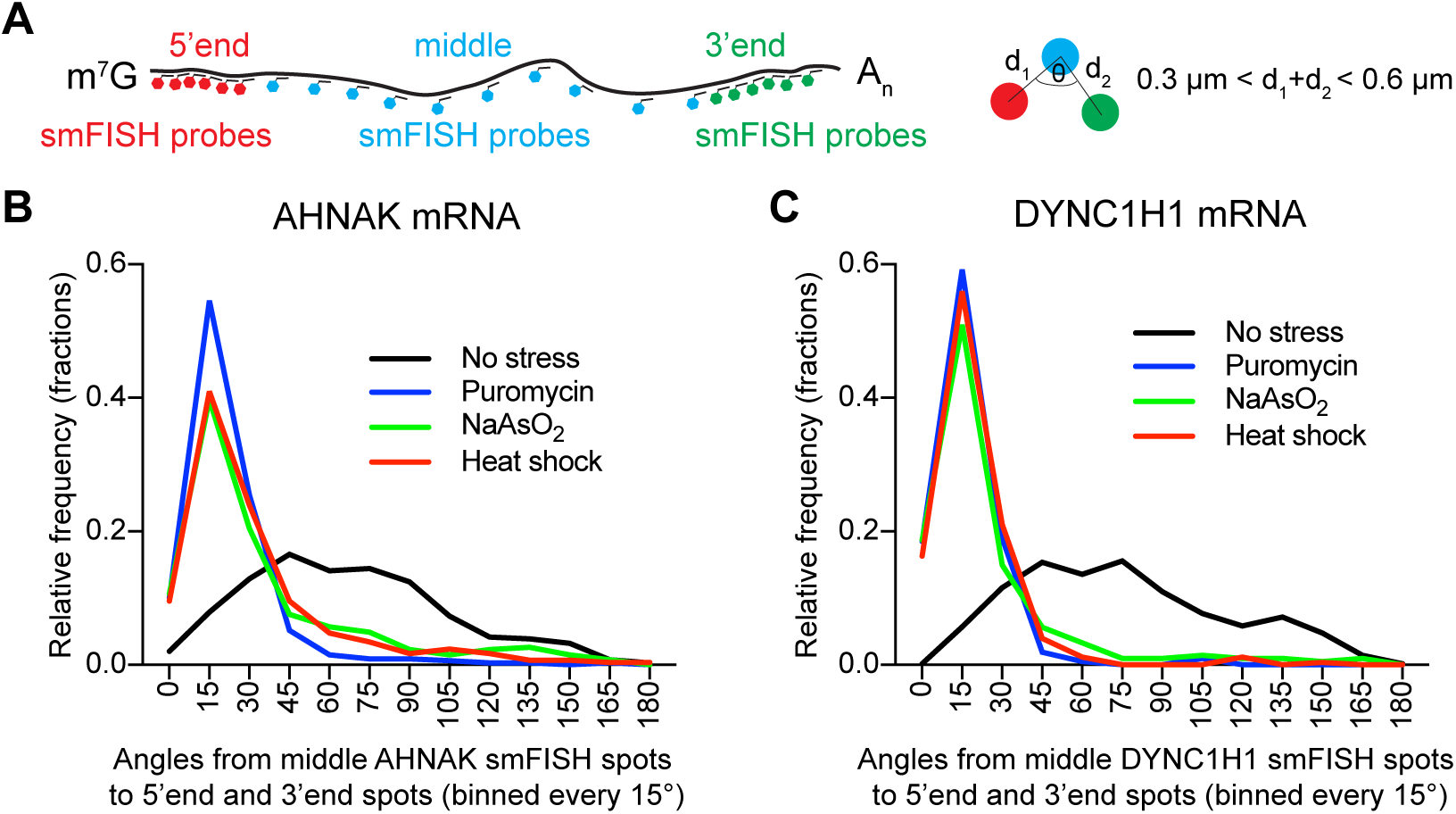
Translation inhibition with puromycin, NaAsO_2_ or heat shock in U-2 OS cells disproportionally shrink the distances between the 5’ and 3’ ends relative the middle of AHNAK and DYNC1H1 mRNPs. **(A)** Cartoon schematic indicating the angles that were measured in **(B, C)**. Analysis was restricted to distances between 0.3 μm to 0.6 μm between 5’-to-middle and middle to 3’-end smFISH spots. Histograms illustrating the relative frequency (fractions) of angles from middle smFISH spots to 5’end and 3’end smFISH spots of **(B)** AHNAK and **(C)** DYNC1H1 mRNAs in unstressed (black line), puromycin-treated (blue), NaAsO_2_-treated (green), or heat shocked (red) U-2 OS cells. The histograms were generated by binning every 15°. More than 200 angles were quantified for each sample.

**Supplemental Figure 6.**
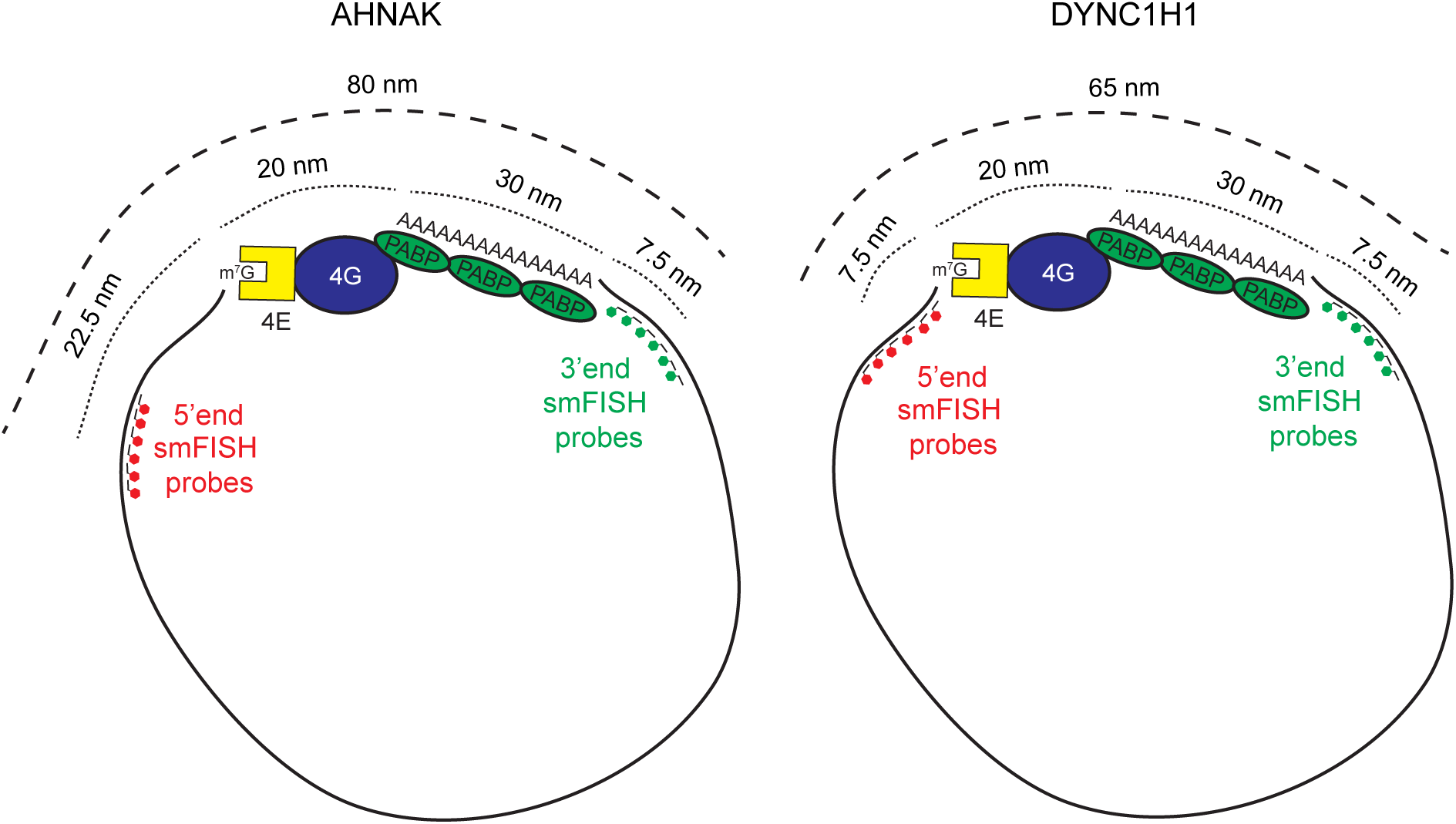
Distances less than 80 nm and 65 nm between 5’ and 3’end smFISH spots for AHNAK and DYNC1H1 mRNAs respectively are consistent with the closed-loop translation model. We estimate the distances between eIF4E, eIF4G, and PABP will be less than 20 nm since an average protein length is 5 nm. With respect to the poly(A) tail length, we estimate the distance between the last nucleotide prior to poly(A) tail and the last A on the poly(A) tail will be 30 nm assuming it is completely extended and the poly(A) tail length is 100 nucleotides. Given that the compaction is ~20 fold relative to contour length, and the contour length of 1000 nucleotides is 300 nm, and given where the 5’end smFISH probes bind on AHNAK and DYNC1H1 mRNAs, we estimate the distance will be approximately 22.5 nm or 7.5 nm from the m^7^G cap respectively. Similarly, we estimate the distance between 3’end smFISH probes and the start of the poly(A) tail for both AHNAK and DYNC1H1 to be 7.5 nm. Therefore, distances less than 80 nm and 65 nm between 5’ and 3’end smFISH spots for AHNAK and DYNC1H1 mRNAs are consistent with the closed-loop model.

